# 53BP1 mediated recruitment of RASSF1A at ribosomal DNA breaks facilitates local ATM signal amplification

**DOI:** 10.1101/2021.11.17.469058

**Authors:** Stavroula Tsaridou, Georgia Velimezi, Frances Willenbrock, Maria Chatzifrangkeskou, Andreas Panagopoulos, Dimitris Karamitros, Vassilis Gorgoulis, Zoi Lygerou, Eric O’Neill, Dafni Eleftheria Pefani

## Abstract

DNA lesions occur across the genome and constitute a threat to cell viability; however, damage at specific genomic loci has a disproportionally greater impact on the overall genome stability. The ribosomal RNA gene repeats (rDNA) are emerging fragile sites due to repetitive nature, clustering and high transcriptional activity. Notably, recent progress in understanding how the rDNA damage response is organized has highlighted the key role of adaptor proteins in the response.

Here we identify that the scaffold and tumor suppressor, RASSF1A is recruited at sites of damage and particularly enriched at rDNA breaks. Employing targeted nucleolar DNA damage, we find that RASSF1A recruitment requires ATM activity and depends on the 53BP1. At sites of damage RASSF1A facilitates local ATM signal establishment and rDNA break repair. RASSF1A silencing, a common epigenetic event during malignant transformation, results in persistent breaks, rDNA copy number alterations and decreased cell viability. Moreover, meta-analysis of a lung adenocarcinoma cohort showed that epigenetic silencing of the scaffold leads in rDNA copy number discrepancies. Overall, we present evidence that RASSF1A acts as a DNA repair factor and offer mechanistic insight in how the nucleolar DNA damage response is organized.

## Introduction

DNA double-strand breaks (DSBs) are the most hazardous lesions arising in the genome of eukaryotic organisms that must be efficiently repaired to secure maintenance of genome integrity and survival. The two main pathways for DSB repair are Non-Homologous End Joining (NHEJ) in which the broken ends are directly ligated and Homologous Recombination (HR) which requires a non-damaged homologous sequence as a template, usually served by the sister chromatid (Gorgoulis et al, 2018; Jackson & Bartek, 2009). The latter process is considered to be error free, however emerging evidence highlights that HR in clustered repetitive loci may be deleterious as it can lead to DNA repeat aberrations and/or chromosomal translocations (Mitrentsi et al, 2020).

The ribosomal RNA repeats (rDNA) that transcribe for the ribosomal RNA are organized in clusters at the Nucleolar Organizer Regions (NORs) at the short arms of the 5 acrocentric human chromosomes. Humans have around 300 rDNA repeats that contain a 13Kb transcribed region and a 30 Kb Intergenic spacer (IGS). The rDNA repeats are transcribed by Polymerase I (Pol I) in a 47S pre-rRNA transcript that then is processed to 18S, 28S and 5.8S rRNAs. Due to recombinogenic instability of the rDNA repeats there is a 10-fold variation in copy numbers among individuals in human populations (Gibbons et al, 2015; Stults et al, 2008). During malignant transformation, replication stress can lead to copy number alterations within the rDNA repeats that have been proposed to serve as a biomarker in therapy treatment or disease severity (Ide et al, 2010; Stults et al, 2009; Wang & Lemos, 2017; Warmerdam et al, 2016). Breaks that arise within the rDNA repeats have been suggested to primarily undergo repair by NHEJ in the nucleolar interior, while persistent breaks relocate to the nucleolar periphery where they get access to the HR machinery (Harding et al, 2015; van Sluis & McStay, 2015; Warmerdam et al, 2016). Break relocation at the periphery serves to separate rDNA that originates from different chromosomes which has been proposed to prevent inter-chromosomal recombination and is driven by nucleolar segregation (Floutsakou et al, 2013; van Sluis & McStay, 2015). These structural changes involve the merge of the fibrillar center (FC) and dense fibrillar component (DFC) of the nucleoli in a bipartite cap-like structure in the nucleolar periphery where rDNA HR repair takes place. Several studies have proposed that nucleolar segregation is driven by ATM dependent transcriptional inhibition in the nucleoli (Harding et al, 2015; Kruhlak et al, 2007; Larsen et al, 2014; Pefani et al, 2018; van Sluis & McStay, 2015), however more recent findings suggest that nucleolar segregation may be transcription independent involving forces arising from Nuclear Envelope invaginations and the actin network (Marnef et al, 2019).

Emerging data show that the recombinogenic nature, high transcriptional activity, formation of secondary structures and clustering of repeats that are localized in different chromosomes constitute the nucleolus a potential hotspot of genomic instability (Lindstrom et al, 2018). Therefore, to avoid the toxic effects of rDNA breaks the nucleolar DNA damage response has evolved certain features including Polymerase I (Pol I) inhibition, dedicated adaptor proteins, chromatin modifications and structural changes to achieve efficient break repair (Korsholm et al, 2019; Kruhlak et al, 2007; Larsen et al, 2014; Mooser et al, 2020; Pefani et al, 2018; van Sluis & McStay, 2015).

RASSF1A is a small tumor suppressor scaffold that we and others have shown to regulate Hippo pathway activity in the presence of genotoxic stress (Pefani & O’Neill, 2016). RASSF1A undergoes frequent promoter methylation early during malignant transformation, which has been linked with early cancer onset and/or worse disease outcome making the scaffold an attractive biomarker (Grawenda & O’Neill, 2015). We previously described the central kinase of the Hippo signaling cascade, MST2, as part of the nucleolar DNA damage response limiting Pol I transcription via phosphorylation of nucleolar chromatin (phosphorylation of H2B at Serine 14) upon double strand break (DSB) formation in rDNA (Pefani et al, 2018). We showed that MST2 activity depends on RASSF1A. RASSF1A is an ATM/ATR target at Serine 131, a phosphorylation that allows dimerization and orientates the associated MST2 monomers to allow stimulation of kinase activity (Hamilton et al, 2009; Pefani et al, 2014). Both RASSF1A and the MST2 kinase are present in the nucleolar fraction independently of damage, however MST2 activation and subsequent H2B phosphorylation depends on ATM activity (Pefani et al, 2018).

Herein we further explore the role of the RASSF1A scaffold at the sites of damage. We identify endogenous RASSF1A localized to DBSs and in particular, a robust recruitment to rDNA. Employing targeted damage at rDNA, we find RASSF1A recruitment to depend on ATM kinase activity and be mediated by 53BP1. 53BP1 is known to promote local ATM signal amplification at break sites, we find RASSF1A to facilitate 53BP1 in local ATM signal establishment at rDNA break sites. RASSF1A downregulation results in compromised local ATM signaling, persistent breaks and decreased cell viability. We propose a model in which RASSF1A acts as a scaffold during initial nucleolar DNA damage response promoting H2BS14 phosphorylation to silence Pol I, via MST2, at the nucleolar interior and subsequently translocates with rDNA breaks to nucleolar caps where ATM signal amplification is required for repair, in a 53BP1-dependent manner. This study provides the first data for direct recruitment of the endogenous scaffold at the sites of damage and offers further insight on how RASSF1A participates in the DNA damage response in a chromatin context.

## Materials and Methods

### Tissue culture and cell treatments

HeLa, U2OS cells were cultured in complete DMEM and a-RPE19 in complete RPMI supplemented with 10% fetal bovine serum in 5% CO_2_ and 20% O_2_ at 37°C. DiVA U2OS cells stably expressing AsiSI-ER-AID (provided by Gaëlle Legube) were cultured in DMEM Glutamax. DSBs in DiVa cells were induced by the addition of 300 nM OHT (Sigma) for 4 h. Cells were transfected with plasmid DNA (2.5 μg/10^6^ cells) or siRNA (50 nM) using Lipofectamine 2000 (Invitrogen) or Lipofectamine RNAiMAX (Invitrogen) according to manufacturer’s instructions. To introduce targeted DSBs in the rDNA IGS gRNA (GATTTCCAGGGACGGCGCCTTGG) was introduced in the pCAS9 vector (OriGene) and transfected in cells. I-PpoI WT and I-PpoI H98A mRNA transfections were conducted as previously described (van Sluis & McStay, 2015). In brief, plasmids were linearised at a NotI site and transcribed using the MEGAscript T7 kit (Ambion). I-PpoI mRNA was subsequently polyadenylated using a Poly(A) tailing kit (Ambion) according to the manufacturer’s instructions. The *in vitro* transcribed mRNA was transfected using the TransMessenger transfection reagent (Qiagen) according to the manufacturer’s instructions. Following 4 h of incubation, the transfection medium was replaced by full medium, and cells were grown for additional 2 h unless stated otherwise. All irradiations were carried out using a Gamma Service^®^ GSRD1 irradiator containing a Cs137 source. The dose rates of the system, as determined by the supplier, were 1.938 Gy/min and 1.233 Gy/min depending on the distance from the source. Cells were exposed in 5 Gy and fixed at the indicated time points. For the laser micro-irradiation experiments, cells seeded on ibidi glass bottom dishes (ibidi μ dish 35mm 81156) were placed in an Olympus Cell-Vivo incubation chamber (37°C, 5% CO_2_) and mounted on the stage of an Olympus IX-83 inverted widefield microscope. Micro-irradiation was performed by using a UV-A pulsed laser (teemphotonics PNV-M02510-355nm-pulse <350psec) coupled to the epifluorescence path of the microscope which was focused to the sample through an Olympus Apochromat 63x/1.2 water immersion objective lens. Operation was assisted by the Rapp Optoelectronics software. Subnuclear irradiations were performed on a 10-μm linear ROI with the use of 3 pulses at 1 % of the total laser power. After the laser micro-irradiation was performed, cells were incubated for 2 h in a normal cell culture incubator (37°C, 5% CO_2_) and were then fixed and stained.

### Drug treatments

The following inhibitors were used: ATMi (KU55933 Selleck, 10 µM), ATRi (VE-821Selleck, 10 µM), Pol Ii (CX-5461 Selleck, 1 µM), Neocarzinostatin (Sigma, 50 ng/µl). Cells were treated with ATM and ATR inhibitors for 1 h prior to I-PpoI mRNA transfections. Cells were treated with CX-5361 for 3 h. Cells were treated with NCS for 30 min and left to recover for 2 h before fixation.

### Immunofluorescence

Cells were grown on coverslips and treated as indicated. Cells were fixed with 4% PFA at RT or with ice cold MEOH at −20°C for 10 min, washed with 1×PBS, permeabilized with 0,5% Triton in 1x PBS and blocked with 2% BSA in 1× PBS. Coverslips were incubated with the indicated antibodies in blocking solution overnight at 4°C, washed and stained with secondary anti-rabbit and or anti-mouse IgG conjugated with Alexa Fluor secondary antibodies (Molecular Probes) for 1h at room temperature. Coverslips were washed with PBS + 0.1% Tween, and DNA was stained with DAPI. For RASSF1A staining the HPA040735 from Atlas Antibodies was used. Cells were analyzed using LSM780 (Carl Zeiss Microscopy Ltd) or Leica SP5 confocal microscopes. Image analysis was done using ImageJ software. *In situ* detection of nascent RNA was performed with the Click-iT Alexa Fluor 488 Imaging Kit (Invitrogen, Molecular Probes) after cells were treated with 0.5 mM 5-EU for 30 min. Cells were analysed using LSM780 (Carl Zeiss Microscopy) or Leica LS5 confocal microscopes, and 5-EU intensity was quantified with the NIS-elements software (Nikon).

### Fluorescent In situ Hybridization

FISH was performed as described in (van Sluis et al, 2016), Briefly, cells were grown in sterile glass coverslips, fixed with 4% PFA, washed with 1x PBS, permeabilized with 0,5% Saponin/0,5% Triton in 1x PBS, washed, incubated with 20% glycerol/PBS for 2h RT and snap frozen in dry ice for 5 min. After thawing in RT, cells were denatured with 0,1 N HCl, washed with 2xSSC and incubated with 50% formamide/2xSSC for 15 min at 37°C. Next, an rDNA probe (human rDNA plasmid pUC-hrDNA-12.0: containing 12 kb that correspond between 30.5-42.5 kb of the rDNA repeat, provided by Brian McStay) diluted in a hybridization buffer (Hybrizol ® VII) was placed in a slide and covered with the coverslip with the cells facing down and sealed with rubber cement. Then, the slide was denatured at 85 °C for 5 min on a heat block and hybridization was carried out at 37 °C for 18 h in an incubator with humidity. Finally, cells were washed three times with 50% formamide/2xSSC at 42 °C, three times with 0,1xSSC (prewarmed at 60 °C) and DNA was stained with DAPI. When FISH was combined with immunofluorescence for UBF (ImmunoFISH), immunofluorescence was performed after FISH as described above. When FISH was combined with immunofluorescence for ATM-pS1981, ATM-pS1981 immunostaining was performed prior to FISH, cells were then fixed with 2%PFA and underwent FISH following 2% PFA fixation.

### Immunoprecipitation

For RASSF1A immunoprecipitation, cells were treated as indicated and washed with ice-cold PBS prior to lysis. Cells were lysed in 1% NP-40 lysis buffer (150 mM NaCl, 20 mM HEPES, 0.5 mM EDTA) containing complete protease and phosphatase inhibitor cocktail (Roche). Total cell extracts were incubated for 3 h with 20 μl protein A Dynabeads (Invitrogen) and 2 μg of RASSF1A antibody (Atlas, HPA040735), HA-tag (HA.C5, Millipore 05-904) or FLAG-tag (M2, Sigma, F3165) at 4°C. Total cell extracts (corresponding to 10% of the immunoprecipitate) and immunoprecipitates were analyzed with western blotting.

### Chromatin fractionation

Following indicated treatments, cells were harvested and the cytosolic was removed by incubation in hypotonic buffer (10 mM HEPES, 10 mM KCl, 1.5 mM MgCl_2_, 0.34 M sucrose, 10% glycerol and 0.1% Triton-X100 supplemented with protease and phosphatase inhibitors) for 10 min on ice following centrifuging. The pelleted nuclei were resuspended in nuclear buffer (10 mM HEPES, 3 mM EDTA, 0.2 mM EGTA, pH 8.0 supplemented with protease and phosphatase inhibitors) and nucleoplasm was released by centrifugation. The final pellet containing the chromatin fraction was resuspended in lysis buffer (10 mM HEPES, 500 mM NaCl, 1 mM EDTA, 1% NP-40, supplemented with protease and phosphatase inhibitor cocktails, Roche, and Benzonase, Millipore), sonicated 3 times at low amplitude, incubated on ice for 15 min and centrifuged to isolate the chromatin fraction. 10 µg of protein from the chromatin fraction were analyzed by western blotting.

### Western Blotting

Extracts were analyzed by SDS–PAGE using a 4-12% Bis–Tris or 3-8% Tris-Acetate NuPAGE gels (Invitrogen) and transferred onto PVDF membranes (Millipore). Subsequent to being washed with PBS containing 1% Tween-20 (PBS-T), the membranes were blocked in 5% milk or 5% bovine serum albumin (BSA) in PBS-T for 1 h at RT and then incubated with the primary antibodies overnight at 4°C. The membranes were incubated with HRP-conjugated secondary antibodies (Cell Signaling) for 1 h at room temperature and exposed to X-ray film (Kodak) or the ChemiDoc Imaging system (Biorad) after incubation with Thermo Scientific Pierce ECL or Clarity ECL (Biorad).

### Quantitatively Real Time PCR

Genomic DNA was extracted using the Nucleospin Genomic DNA from tissue kit (Machenrey-Nagel). Genomic DNA concentration and purity was measured using Nanodrop. rDNA copy number was assessed by quantitative real-time PCR (Applied Biosystems StepOne), using the PowerSYBR® mic (Ambion). rDNA copy number was quantified using the2^ΔΔCt^ method relative to the copy number of the Human GAPDH gene.

### Clonogenic survival assays

Cells were transfected with the indicated siRNAs and 48 h post transfection the appropriate number of cells was seeded in a well of a 6-well plate and treated with (i) the indicated doses of ionizing radiation by a Gamma Service^®^ GSRD1 irradiator or (ii) with 100 nM CX5461 for 5 days. Cells were then grown for a remaining of 7-14 days in regular medium. Plates were then stained with crystal violet (0.5% w/v crystal violet, 50% v/v MeOH and 10% v/v EtOH). For clonogenic survival assays following rDNA DSBs, cells were treated with siRNAs and 48 h post-transfection, I-PpoI WT, or I-PpoI H98A mRNA was introduced with TransMessenger transfection reagent (Qiagen). 6 h post-mRNA transfection, cells were counted, and re-seeded in each well of a 6 well plate. Cells were grown for 10 days in regular medium and stained with crystal violet. Experiments were performed in triplicates. Survival fractions are presented corresponding to relevant viability of each siRNA condition with a DNA damage treatment relative to non-treated cells.

### Antibodies

The following antibodies were used in this study: RASSF1A (HPA040735, Atlas Antibodies), RASSF1A (3F3, Santa Cruz, sc-58470), pS131-RASSF1A (Hamilton et al, 2009), V5 (13202, Cell signalling), γH2AX (JBW301, Millipore, 16-193), γH2AX (2577, Cell Signaling), nucleolin (4E2, ab13541, Abcam), Lamin A/C (4777, Cell Signaling), 53BP1 (NB100-304, Novus Biologicals), 53BP1 (B13, MAB3802, Millipore), MST2 (ab52641, Abcam), Lamin B (ab16048, Abcam), UBF (F9, sc-13125, Santa Cruz), RNF8 (14112-1-A, Proteintech), RIF-1 (A300-569A, Bethyl Laboratories), RPA (Ab2, NA18 Calbiochem), pRPAS33 (A300-246A, Bethyl Laboratories), pRPAS4/8 (Bethyl Laboratories), CHK1(G-4, sc-8408,santa cruz), pCHK1S345 (2348, Cell signaling), CHK2 (C12, 3440), pCHK2Thr68 (9C13C1, cell signaling), SUN1 (ab124770, Abcam), BRCA1 (C9, sc-6954, Santa Cruz), pATMS1981 (GeneTex, GTX40107), ATM (5C2,GTX70107, GeneTex), Treacle (HPA038237, Atlas Antibodies), NBS1 (1D7, GeneTex, GTX70224),NBS1(14956, Cell Signaling) RAD51, (14B4, GTX70230, GeneTex), MRE11 (12D7, ab214 abcam), ATR (E1S3, Cell Signaling), SUN1 (ab124770, Abcam), Fibrillarin (Novus, NBP2-46881).

### Correlation analysis

Methylation data for each patient was downloaded from the GDC portal of the TCGA website (portal.gdc.cancer.gov/projects/TCGA-LUAD). Only data relating to adenocarcinoma were selected, and out of these methylation data was available for 526 patients. The RASSF1A CpG promoter region was defined as the region on chromosome 3 prior to and spanning exon1α. This region is on chr3:50,340,373 - 50,341,109 using GRCh38/hg38 (Malpeli et al, 2019) and contains 85 CpGs, 14 of which are detected using the Illumina 450K array. The mean of the methylation frequencies (the ratio of methylated CpG to total CpG for each site in the dataset) across the 14 sites was used as a readout for RASSF1A methylation. Datasets containing rDNA copy numbers were kindly provided by Dr Wang (Harvard) and were calculated as described in (Wang & Lemos, 2017). All processing of data was performed using R Studio, Version 1.2.5033. Criteria for the selection of cutoff methylation frequencies for each analysis are as described in the results section.

### Statistics

Experiments were performed three times unless stated otherwise. Error bars represent standard deviation. All experiments were performed three times unless otherwise specified in the figure legends. Error bars represent the standard deviation in all figures. Where statistical tests were applied, 75–200 cells were analyzed. For statistical analysis unpaired two-tailed Student’s *t* test was used unless stated otherwise in the figure legend. Error bars represent standard deviation unless stated otherwise in the figure legend. For the rDNA analysis in the LUAD cohort Wilcoxon signed-rank test was used. Statistical significance is depicted with stars (* = 0.05–0.01, ** = 0.01–0.001, *** ≤ 0.001).

### Plasmids

FLAG-RASSF1A and FLAG-RASSF1AS131A were previously described (Hamilton et al, 2009). HA-53BP1WT (aa1-1972), HA-53BP1ΔBRCT (aa 1-1709), HA-53BP1Δ?terminus (aa 921-1972) were previously described (Hansen et al, 2016).

siRNA sequences:

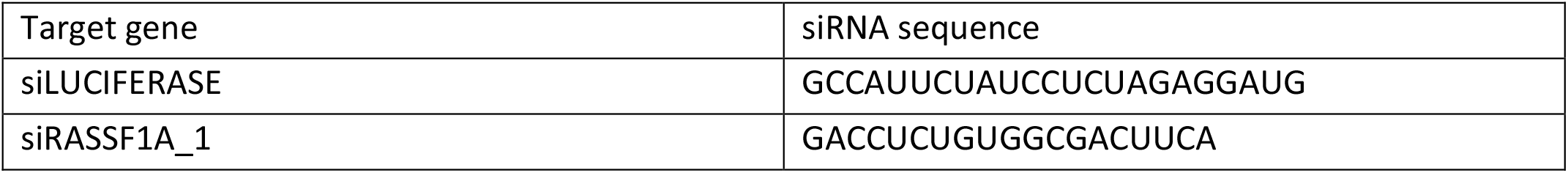
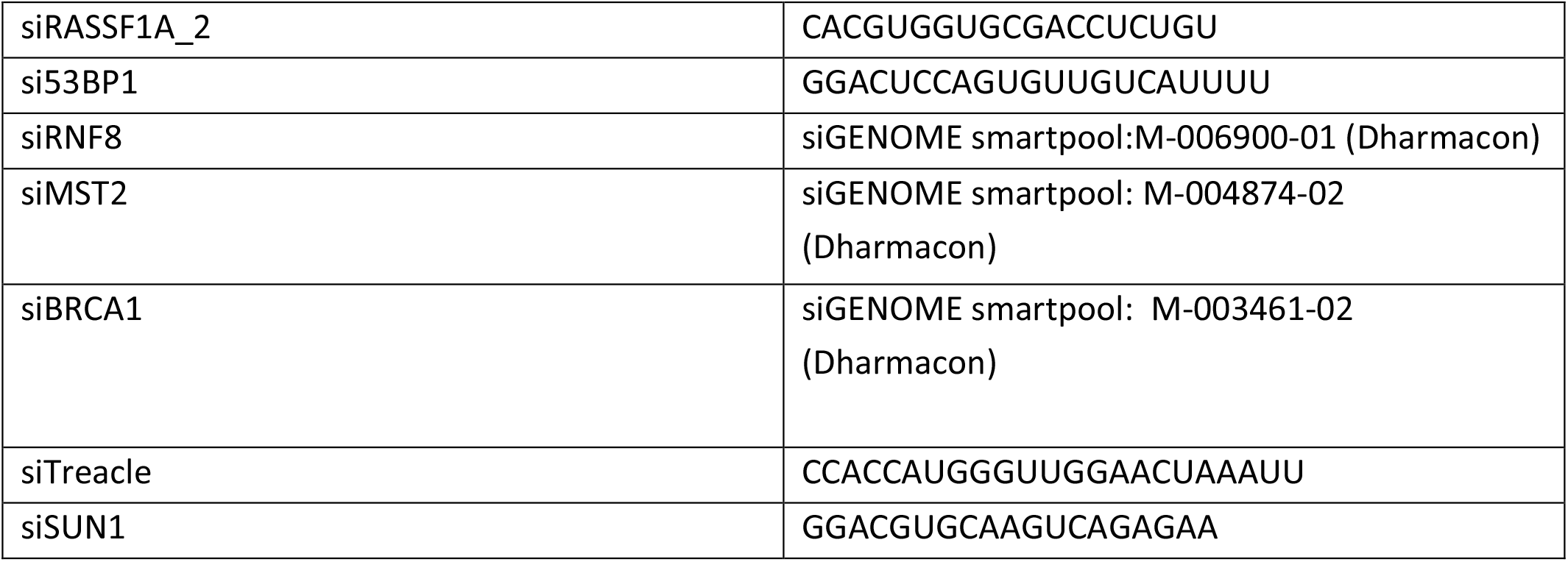

DNA oligos:

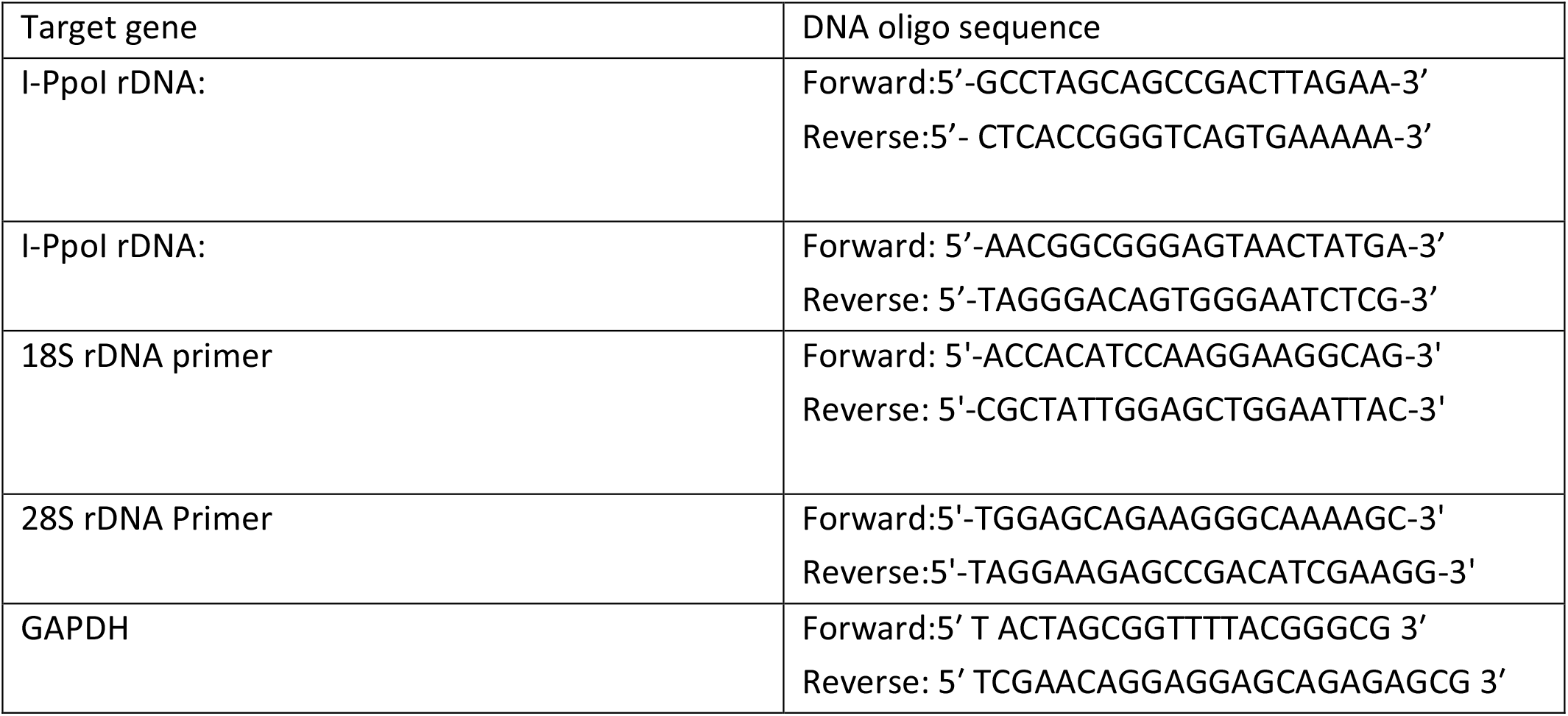

## Results

### RASSF1A is localized at the sites of DNA damage

RASSF1A scaffold is one of the most common epigenetically inactivated genes in human malignancies due to promoter methylation (Dubois et al, 2019; Grawenda & O’Neill, 2015). RASSF1A is a tumor suppressor known to regulate the Hippo signaling cascade in response to genotoxic stress. RASSF1A-mediated activation of MST2 (hippo) promotes stalled replication fork protection (Pefani et al, 2014), apoptosis via the YAP transcriptional co-activator (Hamilton et al, 2009) and regulates Pol I transcriptional activity upon rDNA DSB formation (Pefani et al, 2018). RASSF1A is also involved in repair of UV-induced DNA damage independently of Hippo via interaction with the XPA protein (Donninger et al, 2015). We looked for endogenous RASSF1A localization after exposure to ionizing radiation (γIR) and identified the scaffold co-localizing with γH2AX foci at the sites of damage (Fig 1A, EV1A and EV1C). RASSF1A foci become evident between 30 min and 1 h post irradiation and are no longer detectable 24 h after exposure, when break repair is completed (Fig 1B). To our knowledge this is the first report of the endogenous protein being recruited at the sites of damage. We also observed RASSF1A recruitment at double strand breaks induced by the radiomimetic agent Neocarzinostatin (NCS) and at micro laser generated sites of damage (Fig 1E, EV1F and EV1G). In contrast to UV laser induced damage where the scaffold is recruited throughout the lesion (Fig EV1G), in γIR or NCS induced DNA breaks RASSF1A is located only in a fraction of γH2AX foci (Fig 1A and EV1B), that could be explained by a spatial preference in recruitment. Closer examination of the distribution of RASSF1A foci shows that a significant fraction of RASSF1A lies at nucleoli boundaries where rDNA breaks relocate for HR mediated repair (Fig 1C, 1D, 1E and EV1D, EV1E, EV1F). Moreover, when we measured the distance of RASSF1A foci from fibrillarin marked nucleoli we found that RASSF1A^+ve^/γH2AX^+ve^ foci are located closer to the nucleoli compared to the RASSF1A^-ve^/γH2AX^+ve^ (Fig 1E). To further assess the enrichment of the scaffold in the damaged rDNA loci we used a cell line with inducible expression of the AsiSI endonuclease. AsiSI has several recognition sites within the human genome (Clouaire et al, 2018) and one of those is located at the 5’ EST of the rDNA repeat. Breaks at the 5’ EST of the rDNA repeat induced by AsiSI ectopic expression were recently shown to induce nucleolar segregation and movement of the rDNA breaks to the nucleolar periphery (Marnef et al, 2019). Interestingly, upon AsiSI expression, we observed RASSF1A recruitment mostly at γH2AX foci that are localized at the nucleolar caps indicating a preferential recruitment of the scaffold at the rDNA breaks (Fig 1F and 1G). To get a better insight in whether RASSF1A is involved in rDNA break repair we used the I-PpoI endonuclease to enrich for double strand breaks in the rDNA. As it has been previously described, I-PpoI recognizes a sequence within the 28S-rDNA coding region of each of the approximately 300 rDNA repeats and a limited number of other sites in the human genome (van Sluis & McStay, 2015). In agreement with previous reports, mRNA transfection of V5 tagged I-PpoI results in relocation of rDNA breaks to the nucleolar periphery, formation of γH2AX positive nucleolar caps and downregulation of Polymerase I transcriptional activity, whereas a catalytically inactive version of I-PpoI (H98A) does not result in rDNA DSBs (Fig EV1H, EV1I, EV1J). Indeed, upon induction of rDNA breaks and nucleolar cap formation we observed robust recruitment of RASSF1A scaffold to γH2AX positive nucleolar caps (Fig 1H and 1I). Overall, we find endogenous RASSF1A at a fraction of breaks with evidence supporting robust recruitment at rDNA sites.

**Figure 1.**
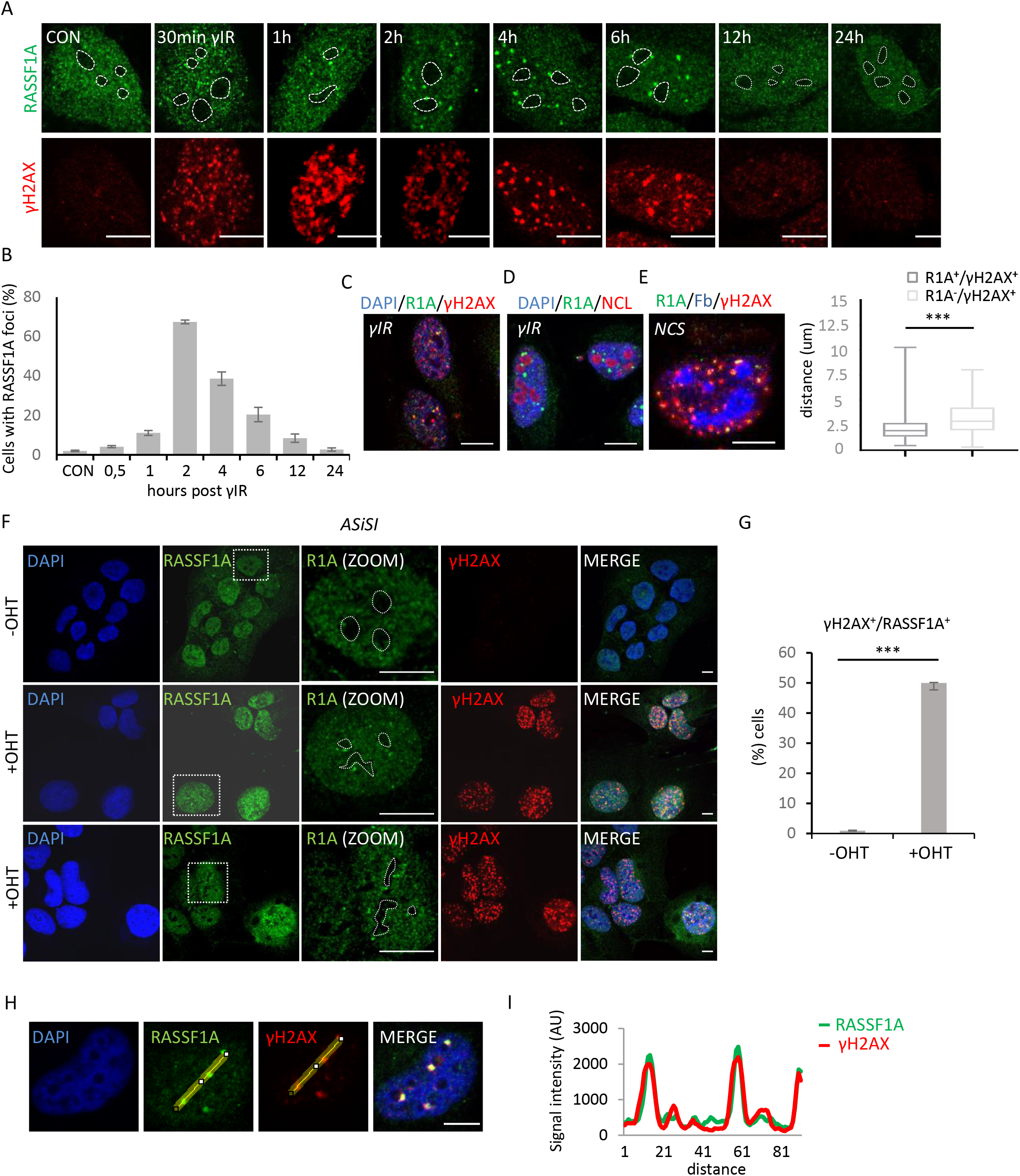
RASSF1A recruitment at double strand breaks. (A) HeLa cells were exposed to 5 Gy ionizing radiation (γIR), collected at the indicated time points and stained for RASSF1A and γH2AX. Images for representative intra-nuclear RASSF1A foci distribution are shown. Nucleolar boundaries are marked with dotted lines. (B) Quantification of the number of cells with RASSF1A foci at the time points presented in (A). Error bars derive from two independent experiments. (C) Cells were treated with 5 Gy γIR and 2 h post treatment fixed and stained for the indicated antibodies. (D) Cells were irradiated with 5 Gy and 2 hours post treatment stained for the indicated antibodies. (E) Cells were treated with 50 ng/ml NCS for 30 min, washed and 2 h later fixed and stained for the indicated antibodies. The distance of RASSF1A^+^/γH2AX^+^ foci or RASSF1A^-^/γH2AX^+^ foci to the closest nucleolus (based on Fibrillarin (Fb) staining) was measured. P value was calculated using a Mann-Whitney test. (F) AsiSI expression in U2OS cells was induced by OHT and cells were stained for RASSF1A and γH2AX. RASSF1A is co-localised with γH2AX foci at the nucleolar caps. Boxed areas are shown in higher magnification. Nucleolar boundaries are marked with dotted lines. (G) Quantification of the number of cells with RASSF1A foci. Error bars derive from three independent experiments. (H) HeLa cells were transfected with I-PpoI endonuclease and stained for RASSF1A and γH2AX. (I) Fluorescence intensity profiles of RASSF1A (green) and γH2AX (red) signals across the HeLa nuclei are shown. Position of line scan indicated by the yellow line. DNA was stained with DAPI. Scale bars = 10 μm.

### RASSF1A recruitment at rDNA DSBs

To acquire a better understanding of RASSF1A temporal recruitment at rDNA breaks we performed a time course after I-PpoI induction with the early marker of rDNA DSBs and upstream regulator of the nucleolar DNA damage response, NBS1 (Korsholm et al, 2019). RASSF1A microfoci are evident from 1 h post I-PpoI mRNA transfection, when nucleolar cap formation starts in a small fraction of cells. Robust RASSF1A staining is observed when nucleolar caps establish between 2 h - 6 h post I-PpoI transfection and RASSF1A foci disappear 24 h post I-PpoI transfection when the majority of breaks have been repaired and cells do not exhibit nucleolar NBS1 staining (Fig 2A and 2B). Similar analysis with other markers known to localize at the rDNA breaks showed that RASSF1A co-localizes with the γH2AX mark and 53BP1 protein at the nucleolar caps and is in proximity with UBF and RPA (Fig EV2A, EV2B and EV2C). We previously showed that there is an endogenous nucleolar fraction of RASSF1A (Pefani et al, 2018), and while formation of microfoci is evident in the nucleolar interior early after induction of rDNA breaks, we only observe robust recruitment at nucleolar caps where breaks relocate for HR-mediated repair (Fig EV2D). When we treated cells with the polymerase I inhibitor CX-5461 that results in rDNA breaks through increasing replication stress in rDNA (Sanij et al, 2020), we similarly observed RASSF1A recruitment to 53BP1^+ve^ nucleolar caps (Fig EV2E). RASSF1A recruitment was also observed in cells with CRISPR/Cas9 induced breaks within the IGS spacer (Fig EV2F), indicating that RASSF1A accumulates at rDNA breaks independent of the break site. To assess potential cell cycle dependency in RASSF1A recruitment at rDNA DSBs we co-stained cells with Cyclin A, a marker of S/G2 phases. RASSF1A is recruited at rDNA DSBs independently of the cell cycle stage (Fig 2C and 2D). We recently reported that there is a fraction of RASSF1A at the nuclear envelope (NE), where it facilitates nucleocytoplasmic actin transport (Chatzifrangkeskou et al, 2019). Methanol fixation facilitates the visualization of NE bound proteins and shows reduced pools of RASSF1A at the NE (LAMIN A/C^+ve^) upon I-PpoI expression indicating that recruitment from the nuclear envelope to the rDNA break sites occurs in response to damage (Fig 2E, 2F and 2G). Taken together this data highlights that RASSF1A is recruited at breaks within the rDNA repeats independently of break site, mechanism of rDNA insult or stage of the cell cycle.

**Figure 2.**
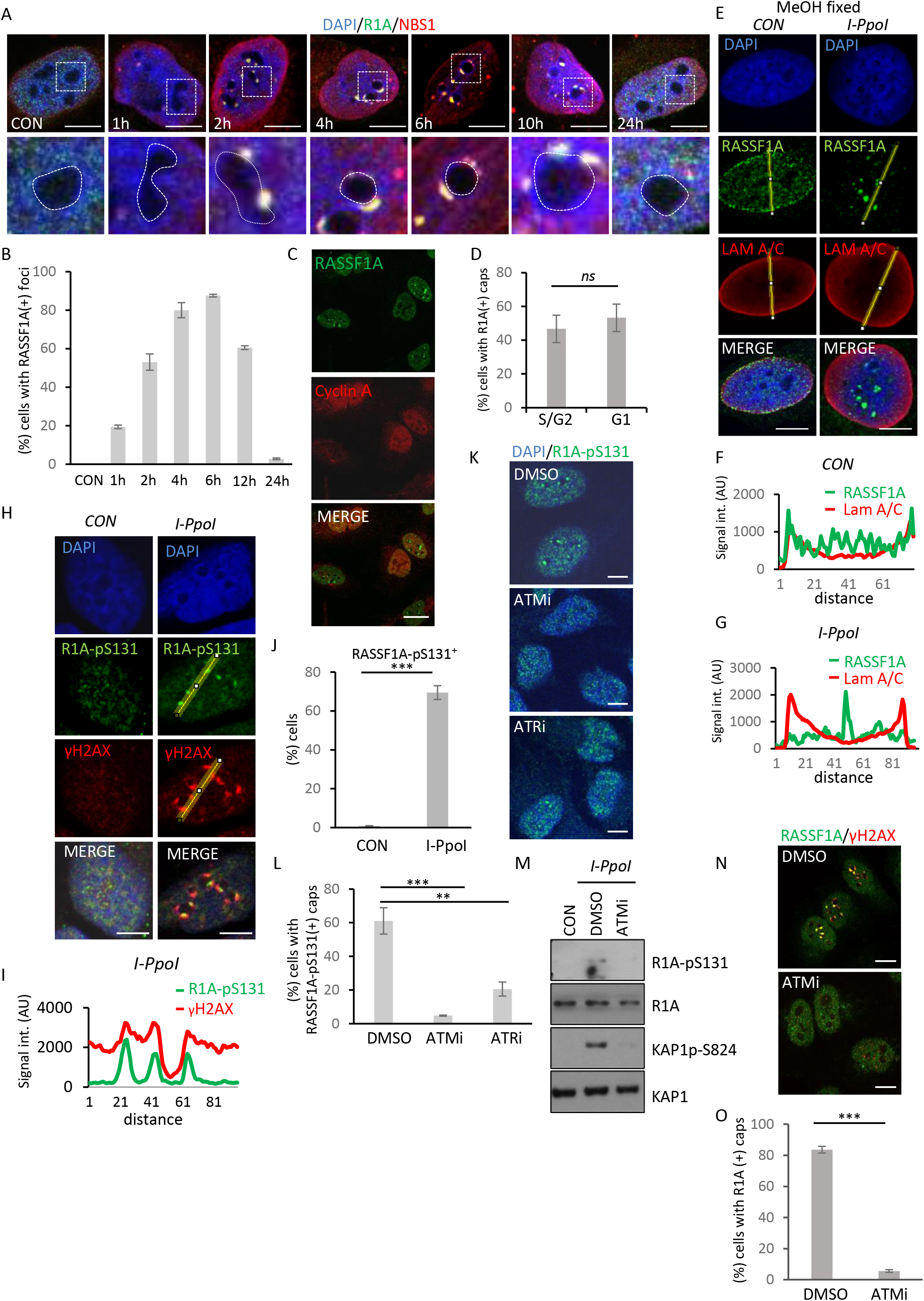
Phosphorylated RASSF1A at Serine 131 localizes at the rDNA DSBs. (A) HeLa cells were transfected with I-PpoI mRNA and collected at the indicated time points. Cells were co-stained for RASSF1A (R1A) and NBS1. Boxed areas are shown in higher magnification. Nucleolar boundaries are marked with dotted lines. (B) Quantification of (A). Error bars derive from two independent experiments. (C) Cells were transfected with I-PpoI mRNA and 6 h post transfection cells were stained for RASSF1A (R1A) and cyclin A. (D) Quantification of (C). Cells were quantified for RASSF1A nucleolar cap recruitment in S/G2 and G1 cells based on Cyclin A staining. Error bars derive from three independent experiments. (E) Assessment of RASSF1A localisation after induction of rDNA DSBs using the I-PpoI endonuclease in MeOH fixed cells. (F, G) Fluorescence intensity profile of RASSF1A (green) and LAMIN A/C (red) signals across the HeLa nuclei in the presence and absence of rDNA DSBs in control (F) and I-PpoI treated cells (G). Position of line scan indicated by the yellow line. (H) Cells were transfected with the I-PpoI mRNA and 6 h later stained for RASSF1A-pS131 and γH2AX. (I) Fluorescence intensity profile of RASSF1A-pS131 (green) and γH2AX (red) signals across the HeLa nuclei are shown. Position of line scan indicated by the yellow line. (J) Quantification of cells with RASSF1A-pS131 positive nucleolar caps 6 h post I-PpoI transfection. Error bars derive from two independent experiments. (K) Cells were treated with the ATMi or ATRi prior to transfection with I-PpoI mRNA. 6 h post mRNA transfection cells were stained for RASSF1A-pS131. (L) Quantification of (K) Error bars derive from two independent experiment. (M) Cells were treated or not with ATM inhibitor (ATMi) followed by I-PpoI mRNA transfection and 6 h later cell lysates were analyzed by western blot for the indicated antibodies. (N) Cells were treated or not with ATMi followed by I-PpoI mRNA transfection and 6 h later cells were fixed and RASSF1A recruitment at rDNA DSBs was assessed. (O) Quantification of (N). Error bars derive from three independent experiments. DNA was stained with DAPI. Scale bars = 10 μm.

### RASSF1A gets phosphorylated at Serine 131 upon rDNA damage induction

ATM has a central role in nucleolar DNA damage response promoting Pol I inhibition, nucleolar segregation, end resection and rDNA DSB repair, while recent studies also showed that the related ATR kinase also contributes to the rDNA break response (Harding et al, 2015; Korsholm et al, 2019; Kruhlak et al, 2007; Mooser et al, 2020; Pefani et al, 2018). Moreover, RASSF1A is a target of both ATM and ATR kinases at Serine 131 (Hamilton et al, 2009; Pefani et al, 2014). Therefore, we looked for RASSF1A phosphorylation at Serine 131 (RASSF1A-pS131) upon rDNA DSB induction. We found that I-PpoI induced damage resulted in increased RASSF1A-pS131 levels and accumulation of the phosphorylated protein at the γH2AX marked rDNA DSB sites (Fig 2H, 2I, 2J and EV3A). Inhibition of ATM or ATR kinases results in decreased establishment of RASSF1A-pS131 at the sites of rDNA breaks (Fig 2K and 2L) with ATM inhibition having a more profound effect. Western blot analysis showed that rDNA damage results in an ATM-dependent increase of RASSF1A-pS131, whilst total RASSF1A protein levels are not affected (Fig 2M). Inhibition of ATM kinase also resulted in abrogation of RASSFS1A foci formation in damaged nucleoli (Fig 2N and 2O). Once again, ATM inhibition had a more profound effect on RASSF1A recruitment compared to inhibition of ATR activity (Fig EV3B and EV3C). In the presence of ATM inhibitor, the RASSF1A NE-associated fraction stays intact, further supporting that the NE-associated fraction of the scaffold is recruited at rDNA DSBs (Fig EV3D). Knock down of Treacle, an upstream adaptor protein that regulates ATR activation at the damaged nucleoli (Mooser et al, 2020), also results in abrogation of RASSF1A recruitment (Fig EV3E and EV3F). Overall, the above data highlight that RASSF1A gets phosphorylated at Serine 131 upon targeted damage in the nucleolus and ATM signaling is required for RASSF1A recruitment at the rDNA damage sites.

### 53BP1 mediates RASSF1A recruitment at rDNA breaks

53BP1 is known to amplify ATM signaling, and within heterochromatic repetitive elements this is important for chromatin decondensation at the sites of damage (DiTullio et al, 2002; Hansen et al, 2016; Lee et al, 2010; Mochan et al, 2003; Noon et al, 2010). Recent observations show that 53BP1 undergoes phase separation to coordinate local DNA damage recognition (Kilic et al, 2019; Pessina et al, 2019). Despite lack of evidence for NHEJ at nucleolar caps we and others observe robust recruitment of 53BP1 at the structures. Given the liquid-like behavior of nucleoli during nucleolar segregation, we reasoned that 53BP1 recruitment at the nucleolar caps could facilitate local ATM signal amplification. We observed 53BP1, its upstream recruitment regulator RNF8 (Schwertman et al, 2016) and partner RIF1 (Chapman et al, 2013) at the nucleolar caps of damaged nucleoli (Fig EV4A). RNF8 DSB associated ubiquitylation activity is necessary for recruitment and retention of 53BP1 and BRCA1 at the DSB sites (Mailand et al, 2007). As expected, depletion of RNF8 results in loss of 53BP1 from nucleolar caps of damaged nucleoli (Fig EV4B and EV4C). Examining the intra-nucleolar cap distribution 53BP1 and other DDR components showed that, like in irradiation induced foci (Chapman et al, 2012; Ochs et al, 2019), there is spatial protein distribution within the nucleolar caps. 53BP1 is in proximity but not colocalizing with RPA, BRCA1 or MRE-11, in agreement with 53BP1 inhibiting end resection (Fig 3A). We observed that RASSF1A follows the same distribution with 53BP1 within the nucleolar caps of damaged nucleoli (Fig 3A and EV4D). We next sought to address whether 53BP1 is involved in the establishment of RASSF1A foci. When 53BP1 or its upstream regulator RNF8 expression is knocked down, RASSF1A recruitment at rDNA breaks was significantly perturbed (Fig 3B, 3C, 3D and 3E). Lack of RASSF1A recruitment is not a consequence of defective segregation, as knockdown of 53BP1 or RNF8 does not affect cap formation (Fig EV4E and EV4F). Given the dependency of 53BP1 recruitment on γH2AX we questioned whether loss of RASSF1A recruitment upon ATM kinase inhibition is attributed to loss of 53BP1 establishment. In agreement with previous observations (Harding et al, 2015) we find robust 53BP1 recruitment at damaged nucleoli upon ATM inhibition potentially due to residual ATR mediated γH2AX establishment (Fig EV4G and EV4H). Knockdown of BRCA1, the recruitment of which also depends on RNF8 (Mailand et al, 2007), did not affect RASSF1A binding at the rDNA DSBs (Fig EV4I and EV4J). Inhibition of ATM signaling or depletion of the 53BP1 adaptor also resulted in reduced RASSF1A foci formation in irradiated cells, indicating that recruitment is independent of the source of the lesion or site of damage (Fig 3F, 3G and EV4K).

**Figure 3.**
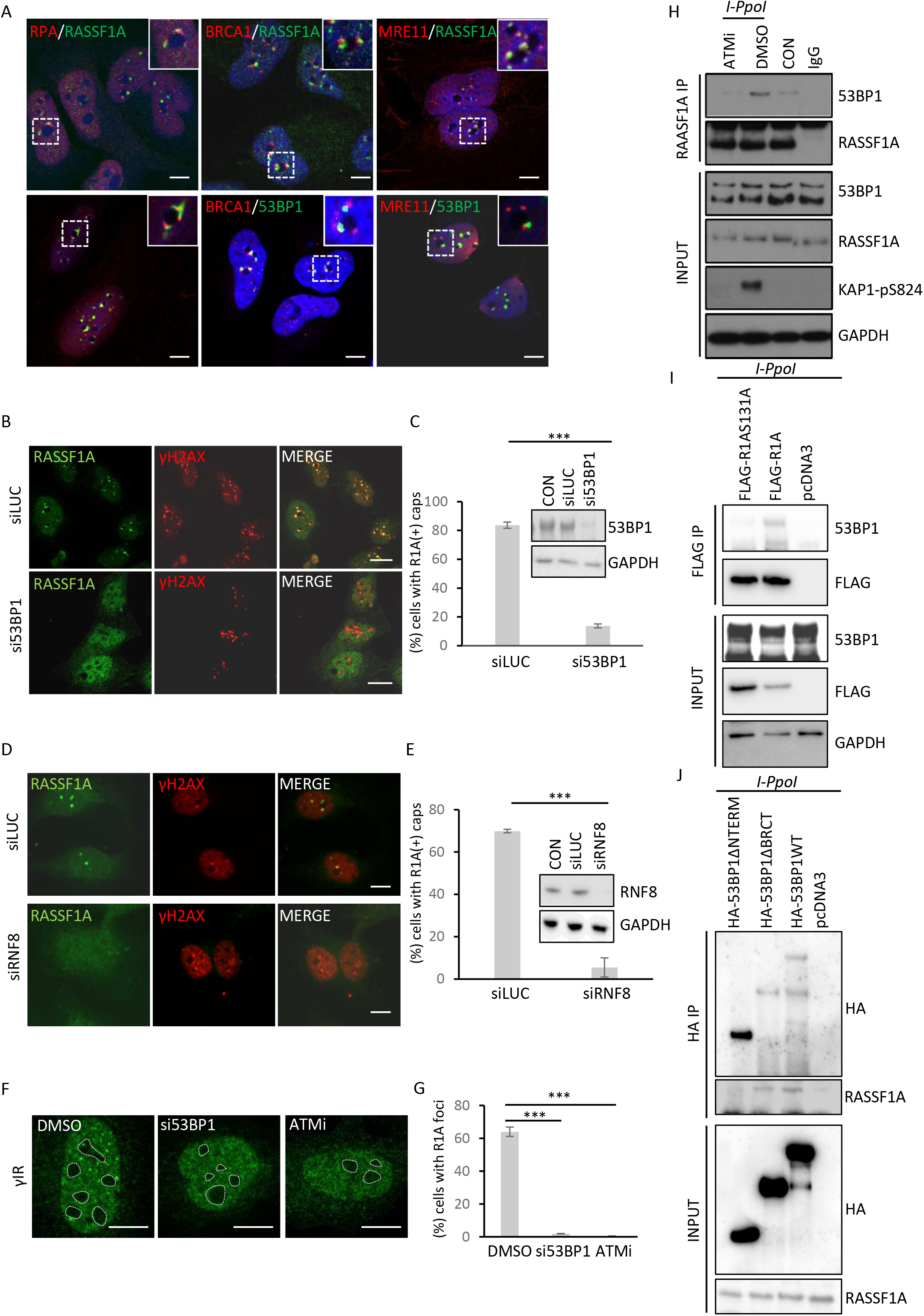
RASSF1A recruitment at rDNA DSBs depends on 53BP1. (A) Cells with I-PpoI induced rDNA breaks were stained with the indicated antibodies. Intra-nucleolar cap localization for RASSF1A (R1A) and 53BP1 in correlation with other proteins was accessed. (B) Cells were treated with siRNAs against LUCIFERASE (LUC) or 53BP1, 48 h later transfected for I-PpoI mRNA, fixed and stained for RASSF1A (R1A) and γH2AX. (C) Quantification of (B) and western blot analysis from siLUCIFERASE (LUC) or si53BP1 treated cells for the indicated antibodies. Error bars derive from three independent experiments. (D) Cells were treated with siRNAs against LUSIFERASE or RNF8, 48 h later transfected for I-PpoI mRNA, fixed and stained for RASSF1A and γH2AX. (E) Quantification of (D) and western blot analysis from siLUCIFERASE (LUC) or siRNF8 treated cells for the indicated antibodies. Error bars derive from three independent experiments. (F) Cells were treated with siRNA against 53BP1, or an ATM inhibitor followed by 5Gy of ionizing radiation. Cells were fixed 2 h post treatment and stained for RASSF1A. (G) Quantification of cells with RASSF1A foci. Error bars derive from two independent experiments. (H) Western blot analysis from total cell extracts with the indicated treatments and RASSF1A immunoprecipitates (IPs) with the indicated antibodies. IgG immunoprecipitation served as a control. (I) Cells were transfected for FLAG-RASSF1A (FLAG-R1A), FLAG-RASSF1A131A (FLAG-R1A131A) or pcDNA3 and 24 h later transfected with the I-PpoI mRNA. Cell lysates were subjected to immunoprecipitation against FLAG. Western blot analysis of the total cell extracts and the IPs is shown. (J) Cells were transfected with either HA-53BP1WT (aa1-1972), HA-53BP1ΔBRCT (aa 1-1709), HA-53BP1Δ?terminus (aa 921-1972) or empty pcDNA3 vector and 24 h later transfected with I-PpoI mRNA. Cell lysates were subjected to Immunoprecipitation for HA. Western blot analysis of the total cell extracts and the IPs is shown. DNA was stained with DAPI. Scale bars = 10 μm.

We next assessed whether 53BP1 is directly responsible for RASSF1A recruitment and indeed, we were able to observe co-immunoprecipitation (IP) (Fig 3H and EV4L) that was significantly induced in response to treatment with ionizing radiation (Fig EV4L) or induction of rDNA DSBs with the I-PpoI endonuclease (Fig 3H). Furthermore, ATM inhibition resulted in decreased RASSF1A-53BP1 interaction (Fig 3H). To examine whether this is a consequence of loss of the ATM-dependent RASSF1A phosphorylation we looked for interaction between 53BP1 with FLAG-tagged RASSF1A or phosphor-site mutant. 53BP1 was significantly reduced in FLAG-RASSF1AS131A IP, suggesting that interaction depends on RASSF1A phosphorylation by ATM (Fig 3I). 53BP1 BRCT domain is not necessary for interaction with RASSF1A, however the N-terminus, known to undergo phosphorylation by ATM and ATR in multiple sites to mediate interaction with PTIP and RIF1 (Mirman & de Lange, 2020), is required for interaction with the RASSF1A scaffold (Fig 3J). Therefore, our data suggests a genotoxic stress induced interaction between 53BP1 and RASSF1A that depends on RASSF1A phosphorylation by ATM and is necessary for recruitment of the scaffold at rDNA DSBs.

### RASSF1A facilitates establishment of the nucleolar DNA damage response

We previously showed that RASSF1A is involved in MST2 activation for nucleolar H2BS14 phosphorylation establishment early in response to induction of damage (Pefani et al, 2018). In agreement with our previous findings, we observed increased nucleolar 5-EU incorporation in RASSF1A knock down cells (Fig EV5A, EV5B and EV5C). Given that RASSF1A is necessary for nucleolar MST2 activity towards H2BS14 phosphorylation to facilitate Pol I transcriptional inhibition (Pefani et al, 2018), we looked at whether RASSF1A depletion affects nucleolar segregation. Several studies have highlighted the link between transcriptional inhibition and nucleolar segregation, however recent data suggests that the two processes may be uncoupled (Fages et al, 2020; Marnef et al, 2019). siRASSF1A cells show a decrease in fully segregated nucleoli, whilst most of the cells presented a partially segregated phenotype with formation of UBF condensates that do not fully move to the nucleolar exterior (Fig EV5D and EV5E). Limited segregation was also reported in siMST2 cells, in agreement with RASSF1A-MST2 mediated Pol I transcriptional regulation being involved in nucleolar segregation (Pefani et al, 2018). Depletion of RNF8 or 53BP1 does not result in nucleolar segregation defects (Fig EV4E and EV4F), suggesting that RASSF1A recruitment at rDNA DSBs which is compromised in 53BP1 knock down cells is not involved in regulation of nucleolar segregation.

53BP1 was previously shown to act in local ATM signal amplification and facilitate phosphorylation of ATM substrates upon exposure to γIR (DiTullio et al, 2002; Lee et al, 2010; Noon et al, 2010). To examine whether RASSF1A has a direct role in rDNA break repair, we knocked down RASSF1A and examined the establishment of the nucleolar DNA damage response (Fig 4A-D). We therefore looked for ATM signal establishment (Fig 4A and 4B), ssDNA formation (Fig 4C) and homology mediated repair (Fig 4D) in cells transfected with I-PpoI mRNA. ATM-pS1981 and the downstream target KAP1-pS824 were reduced at the nucleolar caps of siRASSF1A treated cells (Fig 4A and 4B), indicative of a defective spatial concentration of ATM activity similar to that observed upon 53BP1 depletion (Fig 4E and 4F). Given the partial segregation phenotype of RASSF1A depleted cells (Fig EV5D and EV5E) we wanted to further address whether lack of ATM signal establishment is due to lack of rDNA break movement to nucleolar periphery (Fig 4G and 4H). In agreement with unperturbed formation of nucleolar caps in si53BP1 treated cells (Fig EV4E and EV4F) we observed robust mobilization of rDNA in the nucleolar periphery (Fig 4H). rDNA mobilization was also evident in siRASSF1A treated cells despite defective nucleolar segregation based on UBF staining (Fig 4H). Moreover, in ImmunoFISH experiments where cells were stained for rDNA and ATM-pS1981 we observed lack of active nucleolar ATM despite rDNA exposure at the nucleolar exterior (Fig 4G).

**Figure 4.**
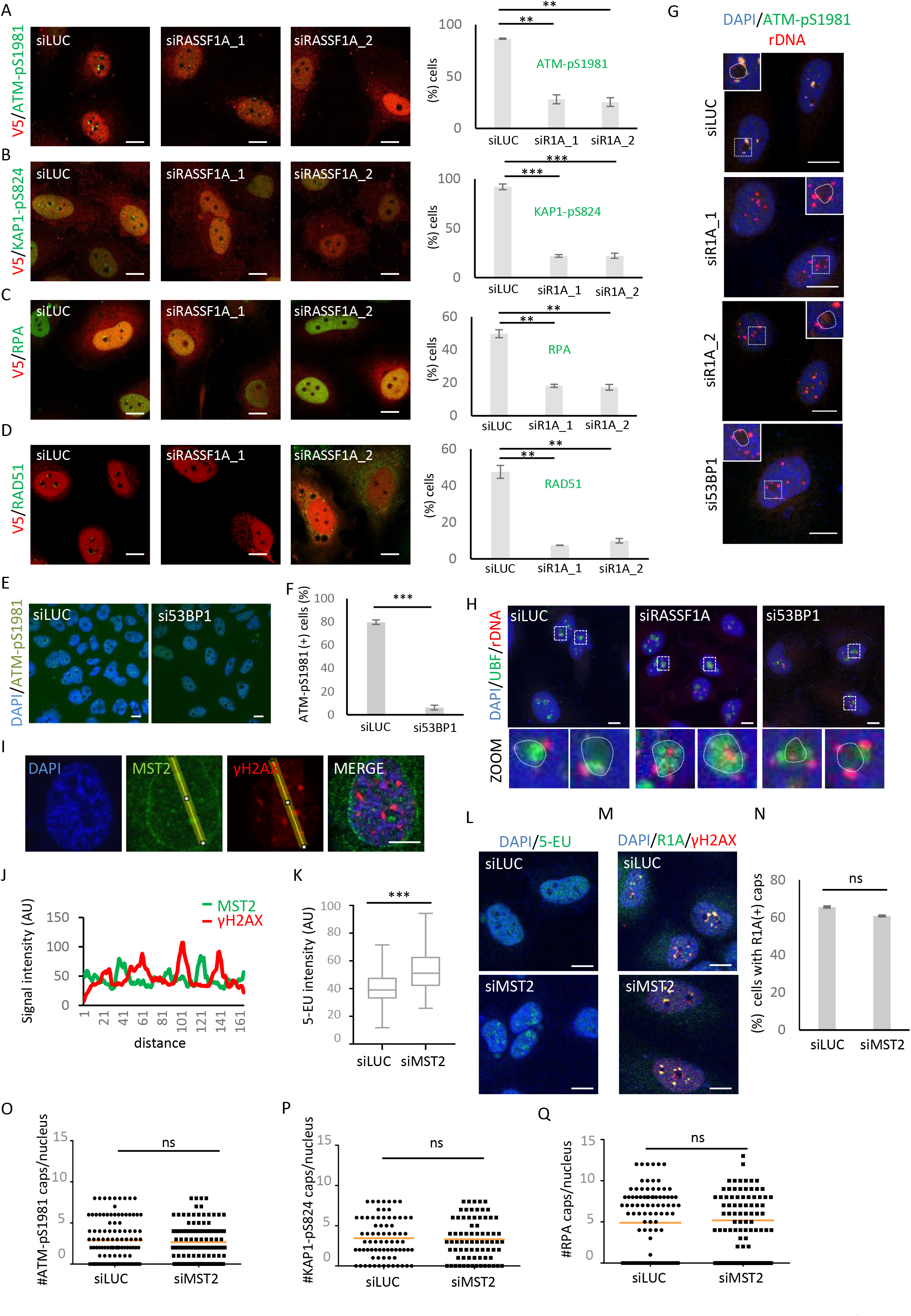
RASSF1A depletion results in impaired rDNA damage response. (A-D) HeLa cells were treated with siLUCIFERASE (siLUC) or two different siRNAs against RASSF1A (siR1A_1 and siR1A_2) and 48 h later cells were transfected with I-PpoI mRNA to induce rDNA DSBs. 6 h post mRNA transfection, cells were stained for V5 to identify I-PpoI transfected cells and the other indicated antibodies. Representative images and quantification of staining for the indicated antibodies are shown. Error bars derive from three independent experiments. (E) Cells were treated with siLUCIFERASE (siLUC) or si53BP1 and 48 h later transfected with I-PpoI mRNA and stained for ATM-pS1981. (F) Quantification of (E). Error bars derive from three independent experiments. (G) Cells were treated with siLUCIFERASE (siLUC), siRASSF1A or si53BP1 and 48 h later transfected with the I-PpoI mRNA. Cells were fixed and underwent ImmunoFISH against ATM-pS1981 and an rDNA probe. Boxed areas are shown in higher magnification. Nucleolar boundaries are marked with dotted lines. (H) Cells were treated with siLUCIFERASE (siLUC), siRASSF1A or si53BP1 and 48 h later transfected with the I-PpoI mRNA. Cells were fixed and underwent ImmunoFISH against UBF and an rDNA probe. Boxed areas are shown in higher magnification. Nucleolar boundaries are marked with dotted lines. (I) HeLa cells were transfected with I-PpoI endonuclease and stained for MST2 and γH2AX. (J) Fluorescence intensity profiles of MST2 (green) and γH2AX (red) signals across the HeLa nuclei are shown. Position of line scan indicated by the yellow line. (K) Quantification of 5-EU intensity in siLUCIFERASE (siLUC) or siMST2 treated cells transfected with I-PpoI mRNA. Error bars derive from two independent experiments. P values were calculated using a Mann-Whitney test (L) Representative images of cells quantified in (K). Cells were treated with 5-EU for 30 min prior to fixation and assessed for incorporation.. (M) Cells were treated with siLUCIFERASE (siLUC) or siMST2 and 48 h post transfection cells were transfected with I-PpoI mRNA. 6 h post transfection cells were fixed and stained for RASSF1A (R1A) and γH2AX. (N) Quantification of (M). Error bars derive from three independent experiments. (O-Q) Quantification of ATM-pS1981 (O) KAP1-pS824 (P) and RPA (Q) in cells treated with siLUCIFERASE (siLUC) or siMST2 and 48 h later transfected with I-PpoI mRNA. Data derives from two independent experiments. P values were calculated using a Mann-Whitney test. DNA was stained with DAPI. Scale bars = 10 μm.

RPA foci formation is commonly used as a proxy for ssDNA accumulation. End resection based on RPA foci formation in damaged rDNA, in contrast to other genomic loci, is placed downstream of ATM/ATR signaling (Fig EV5F and EV5G) (Korsholm et al, 2019; Mooser et al, 2020). RASSF1A knockdown phenocopies the effect of ATM inhibition on the establishment of RPA foci, in agreement with defective nucleolar ATM signal establishment in cells knock down for RASSF1A (Fig 5C). Moreover, reduced RPA phosphorylation (RPA-pS4/8 and RPA-pS33) was observed in I-PpoI treated cells knocked down for RASSF1A (Fig EV5H). Compromised local ATM activation upon RASSF1A depletion, also results in perturbed homology mediated repair marked by decreased recruitment of RAD51 at the nucleolar caps (Fig 5D). We previously reported that RASSF1A indirectly protects stalled replication forks from extended resection via BRCA2 stabilization of RAD51 foci (Pefani et al, 2014), this function of RASSF1A could also result in reduced RPA/RAD51 staining. RASSF1A downregulation does not affect 53BP1 nucleolar cap recruitment in agreement with the scaffold acting downstream 53BP1 (Fig EV5I and EV5J).

**Figure 5.**
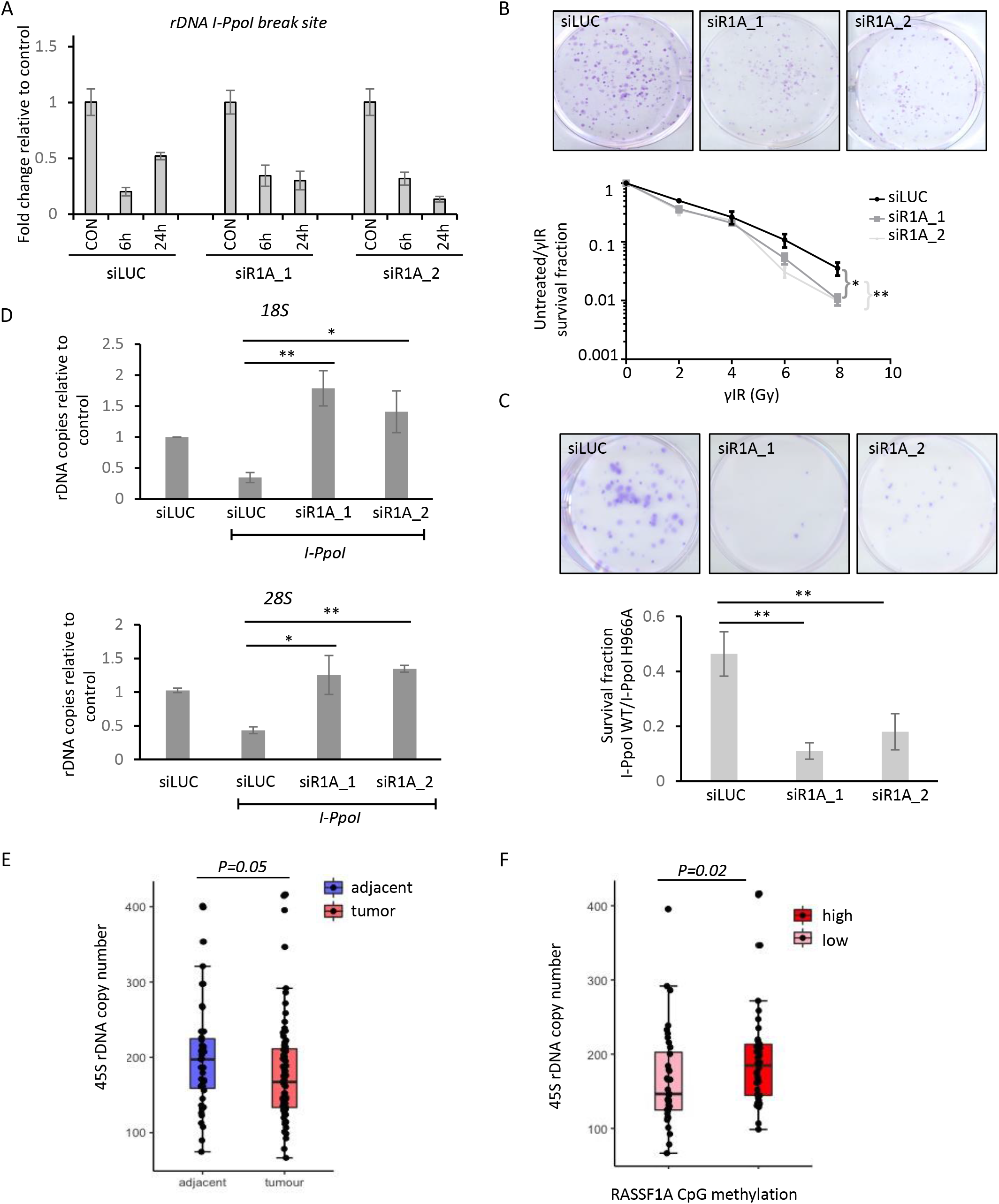
RASSF1A deletion results in reduced cell viability and rDNA copy number alterations. (A) HeLa cells were treated with the indicated siRNAs following treatment with I-PpoI mRNA to induce rDNA DSBs. Genomic DNA was isolated at the indicated times post mRNA transfection and quantitative real time PCR was performed with primers spanning the I-PpoI recognition site. Abundancy of rDNA copies relative to control cells was quantified in each siRNA condition. Error bars derive from two independent experiments. (?) Clonogenic survival analysis of HeLa cells treated with the indicated siRNAs and the indicated doses of Ionizing radiation (γIR). Survival fraction was calculated. Error bars represent SEM and derive from two independent experiments. P-values were calculated with two way-Anova. (C) Clonogenic survival analysis and representative images of HeLa cells treated with the indicated siRNAs and I-PpoI WT. Survival ratio I-PpoI WT/I-PpoI H98A in each siRNA condition is presented. Error bars derive from three independent experiments. (D) HeLa cells were treated with the indicated siRNAs and transfected or not with I-PpoI mRNA. 96 h post mRNA transfection genomic DNA was isolated and the rDNA copy number in the I-PpoI WT transfected cells relative to I-PpoI H98A transfected cells was accessed by real time PCR. Error bars derive from two independent experiments. (E) 45S rDNA copy number analysis in patients’ tumors compared to adjacent controls in the LUAD cohort. (F) 45S rDNA copy number analysis in the LUAD patient samples based on RASSF1A CpG promoter methylation.

To understand whether regulation of nucleolar MST2 kinase activity is involved in RASSF1A recruitment at nucleolar caps or the subsequent establishment of local ATM activation, we first looked for MST2 kinase localization at damaged nucleoli. Consistently with our previous observations for H2B-pS14 intranucleolar localization (Pefani et al, 2018), MST2 is found at the nucleolar interior of damaged nucleoli and not at the nucleolar caps where RASSF1A relocates together we persistent breaks (Fig 4I and 4J). MST2 knockdown (Fig EV5K) results in continued 5-EU incorporation in damaged nucleoli as previously described (Fig 4L and 4K), however under the same depletion conditions we did not observe any effect on RASSF1A nucleolar cap localization (Fig 4M and 4N), ATM-pS1981 (Fig 40 and EV5L), KAP1-pS824 (Fig 4P and EV5M) or RPA (Fig 4Q and EV5N) establishment at nucleolar caps. The above data suggests that RASSF1A function at the sites of rDNA breaks does not involve activation of the MST2 kinase.

### RASSF1A depletion results in persistent breaks, rDNA copy number aberrations and reduced survival

To assess the impact of impaired ATM signal establishment at damaged nucleoli upon decreased RASSF1A expression we monitored rDNA break repair kinetics at the rDNA break sites. Indeed, RASSF1A knock down cells retain a higher number of rDNA breaks 24 h following I-PpoI induction, indicative of defective repair (Fig 5A and EV6A). To examine whether persistent breaks observed in the absence of RASSF1A could affect cell survival, we studied radiosensitivity of RASSF1A knocked down cells. Cell viability was affected in a γIR dose dependent manner in cells with reduced RASSF1A expression (Fig 5B). Previous studies have highlighted the toxicity of rDNA breaks and their impact in cell survival (Warmerdam et al, 2016). To examine whether loss of RASSF1A also affects cell viability in the context of enriched DNA damage in the nucleolus we performed clonogenic survival assays upon I-PpoI induced rDNA break formation. RASSF1A knock down resulted in significant reduction of cell viability in agreement with a role of the scaffold in promoting rDNA break repair (Fig 5C). Similar results were also obtained when knock down of RASSF1A was tested in combination with the CX-5461 Pol I inhibitor that results in rDNA break formation (Fig EV6B).

Due to the highly repetitive nature of the 45S rDNA arrays, breaks within the repeats may lead to excessive recombination and aberrations in the number of rDNA repeats. It was previously reported that rDNA breaks induced by I-PpoI result in loss of rDNA repeats in an HR dependent manner (Warmerdam et al, 2016). We performed qPCR analysis on genomic DNA isolated from cells that had undergone rDNA damage and were treated with control siRNA or siRNAs against RASSF1A. We noticed that in contrast to controls that show decreased rDNA copies, surviving cells with low levels of RASSF1A did not exhibit loss of rDNA copies following induction of rDNA DSBs (Fig 5D). Similar results were obtained for cells depleted for BRCA1, a key HR protein (Prakash et al, 2015) (Fig EV6C).

A recent study by Wang and Lemos in patient cohorts showed that in several cancers 45S rDNA repeats are lost as a result of recombinogenic events due to accumulation of genomic instability (Wang & Lemos, 2017). Epigenetic loss of RASSF1A expression via promoter methylation is a common early event in lung malignant transformation (Grawenda & O’Neill, 2015). To address if RASSF1A promoter methylation affects 45S rDNA copy number, we explored publicly available data from the lung adenocarcinoma cohort (LUAD) of The Cancer Genome Atlas from which we extracted tumor information with *RASSF1A* promoter CpG island methylation status using Genomic Data Commons Data portal. Previous analysis showed that LUAD tumor samples presented fewer copies of 45S rDNA repeats compared to adjacent tissue (Wang & Lemos, 2017). We similarly observed 45S repeat loss in the fraction of LUAD samples for which we had information on both RASSF1A promoter methylation status and 45S rDNA copy number (tumor=74 /adjacent=42) (Figure 5E). We then sought to address whether RASSF1A promoter methylation affects 45S rDNA copy number within the tumor samples. We analyzed the fractional methylation of a total 463 samples from the LUAD cohort and 63 samples of adjacent tissue, for which RASSF1A promoter methylation data was available, to set a cutoff for «high» and «low» methylation samples (Fig EV6D). «Highly» methylated are considered the samples that cluster higher than normal tissue RASSF1A promoter methylation levels. Tumor samples with fractional RASSF1A promoter methylation > 0.198 are listed as «highly methylated» and tumor samples with fractional methylation < 0.12 are listed as «low methylated». We observed higher number of 45S rDNA repeats in the RASSF1A highly methylated tumors compared to the low methylated samples (Fig 5F), indicative of retention of rDNA copies in those tumors. This is in agreement with the hypothesis that RASSF1A facilitates Homologous-mediated repair, a driver of rDNA copy number loss (6, 7). We hypothesize that cells that have lost RASSF1A expression mostly rely on NHEJ for repair that takes place at the nucleolar interior (Harding et al, 2015).

Taken together the above data identify RASSF1A scaffold as a DNA repair factor that localizes at DSBs. We find RASSF1A in a subset of breaks and robust recruitment at rDNA break sites. To our knowledge this is the first report of endogenous RASSF1A localizing at the sites of DNA damage. rDNA loci are emerging fragile sites, and the nucleolar DNA damage response aims to secure efficient repair with reduced occurrence of translocations due to clustering of the repeats. We found that RASSF1A recruitment at rDNA sites depends on ATM activity and interaction with 53BP1. RASSF1A is necessary for MST2 kinase activation at the nucleolar interior for subsequent downregulation of Pol I transcription (Pefani et al, 2018). MST2 kinase does not relocate to the nucleolar caps, does not affect RASSF1A recruitment at the sites of damage and does not phenocopy perturbed establishment of ATM signaling. Therefore, we propose a model in which RASSF1A acts as a multifunctional adaptor during rDNA break repair via regulation of MST2 activity to establish H2BS14 phosphorylation in the nucleolar interior (Pefani et al, 2018), and recruitment at persistent rDNA breaks in a 53BP1 dependent manner to facilitate local ATM signal establishment for efficient break repair (Fig 6). RASSF1A epigenetic inactivation, a process often observed during malignant transformation, results in failure to establish a local DDR, persistent breaks and increased genomic instability further supporting the role of the scaffold as a tumor suppressor.

**Figure 6.**
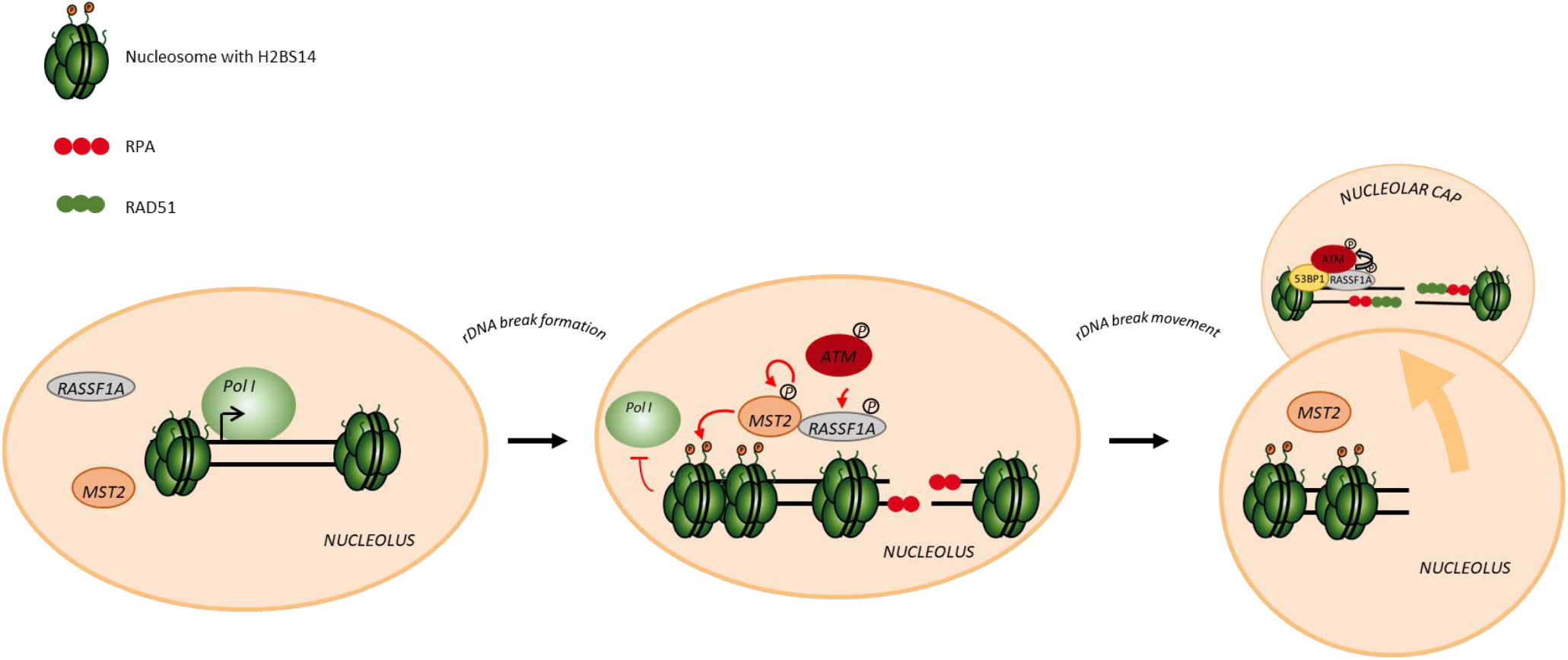
Model for the role of the RASSF1A scaffold in rDNA repair. RASSF1A is a multifunctional protein that acts in the nucleolar DNA damage response via regulation of (1) nucleolar chromatin dynamics in an MST2 kinase dependent manner in the nucleolar interior and (2) rDNA break repair in a 53BP1 dependent manner at the nucleolar caps (see text for details).

## Discussion

In this study we provide evidence that RASSF1A adaptor acts as a bona fide DNA repair factor that accumulates at DSBs. We and others have previously shown the involvement of the scaffold in maintenance of genome stability via or independently of the Hippo cascade (Donninger et al, 2015; Pefani et al, 2014). However, this is the first study to identify endogenous RASSF1A at the break sites colocalizing with DNA DSB markers. We find RASSF1A in a subset of γH2AX foci induced by γIR or radiomimetic agents and robust recruitment in the nucleolar periphery, where rDNA breaks relocate for homology mediated repair. Despite enrichment, a fraction of the DNA repair RASSF1A^+ve^ foci are not linked with the nucleoli, indicating that RASSF1A also marks other sites of damage. In future studies it would be interesting to examine whether these sites could be non-active rDNA repeats that localize outside the nucleoli and have been recently reported (Potapova et al, 2019), or include other repetitive elements (e.g. telomeric or centromeric repeats). The preference in RASSF1A binding could be provided via the chromatin environment or protein interactions (Jeggo et al, 2017).

Given the emerging role of rDNA as a hub of genomic instability (Lindstrom et al, 2018) and our previous findings that place the scaffold as part of the nucleolar DNA damage response (Pefani et al, 2018) we focused on understanding the significance of the recruitment of the scaffold at rDNA sites by employing targeted DNA damage. We previously showed that RASSF1A is involved in the nucleolar DNA damage response via regulation of MST2 kinase activity that phosphorylates nucleolar H2B at Serine 14 facilitating Pol I transcriptional repression. In cell fractionation experiments RASSF1A was found in the nucleolus independent of the presence of damage (Pefani et al, 2018). Interestingly, neither MST2 nor H2B-pS14 were localized at the γH2AX^+ve^ caps of the damaged nucleoli, although they are both found in the interior (Pefani et al, 2018). H2B-pS14 could mark either rDNA repeats that do not move to the periphery upon nucleolar segregation or evicted histones that have been released in the nucleolar interior to allow repair (Hauer & Gasser, 2017). In this study, we identify a fraction of RASSF1A at rDNA DSBs that have re-located to the nucleolar exterior. Further analysis showed that RASSF1A gets phosphorylated upon rDNA DSB formation by ATM at Serine 131 and ATM signaling is required for recruitment at the rDNA breaks.

53BP1 adaptor is mostly studied as a resection inhibitory factor that promotes NHEJ during G1 phase of the cell cycle (Zlotorynski, 2018). 53BP1 has also an established, but less well understood, role in promoting local ATM signal amplification mostly studied within repetitive heterochromatic loci, where ATM mediated phosphorylation of KAP1 at Serine 824 triggers chromatin relaxation (Lee et al, 2010; Mochan et al, 2003; Noon et al, 2010). ATM activity at the γIR foci was shown to be stimulated by 53BP1 mediated interactions with MRN (Lee et al, 2010). Recent findings highlight that 53BP1 undergoes phase separation to integrate DNA repair factors to organize large repair compartments (Kilic et al, 2019). Liquid phase separation events are employed to achieve heterochromatin compartmentalization for break repair (Rawal et al, 2019) and are central in nucleolar organization (Feric et al, 2016). 53BP1 could have a central role in organizing DNA repair at genetic loci where break movement is important to achieve repair. We and others have found that inhibition of ATM signaling does not significantly affect 53BP1 establishment at the nucleolar exterior (Harding et al, 2015) potentially due to adequate γH2AX establishment via ATR. Knockdown of 53BP1 or its recruitment regulator RNF8 resulted in impaired establishment of RASSF1A without affecting nucleolar cap formation. Interaction between RASSF1A and 53BP1 requires ATM signaling as the RASSF1AS131A mutant showed reduced ability to bind to 53BP1. Moreover a 53BP1 N-terminal deletion mutant that lacks the heavily ATM/ATR phosphorylated sites (Mirman & de Lange, 2020) is also necessary for interaction between the two adaptors. Defective recruitment of the scaffold impacts on local ATM signaling establishment, a phenotype observed upon depletion of the 53BP1 adaptor but not in response to MST2 knock down. 5-EU incorporation is significantly increased in siMST2 treated cells, suggesting that lack of any ATM signal establishment phenotype is not due to residual kinase activity. Staining for RAD51 showed that cells that present low levels of RASSF1A have reduced stability of the recombinase, a phenotype that was also previously reported at stalled replication forks of *RASSF1A* knock out cells (Pefani et al, 2014).

ATM localization and signal amplification are important for efficient nucleolar DNA damage response and other adaptor proteins including Treacle, TopBP1 and NBS1, have been shown to participate in ATM/ATR signal establishment (Korsholm et al, 2019; Larsen et al, 2014; Mooser et al, 2020; Noon et al, 2010). Therefore, we propose a model in which induction of rDNA DSBs results in activation of pre-existing nucleolar pools of RASSF1A and MST2 in the nucleolar interior, promoting H2BS14 phosphorylation (Pefani et al, 2018). Following movement of persistent breaks in the nucleolar periphery and formation of DNA repair protein clusters at the nucleolar exterior, RASSF1A is recruited by 53BP1 at nucleolar caps where it facilitates nucleolar signal establishment (Fig 6). At the nucleolar caps due to condensate formation and break clustering RASSF1A signal is robust while in the nucleolar interior potentially remains diffused. Whether pools from nucleolar interior or only other fractions (i.e. the Nuclear Envelope as shown here) contribute to nucleolar cap recruitment is still not clear.

Translocation of the rDNA breaks has been proposed to serve in separating each NOR in a distinct nucleolar cap (Floutsakou et al, 2013). Break repositioning for homology mediated repair has been also observed in heterochromatic elements indicating that break mobilization is linked with repetitive element clustering (Chiolo et al, 2011; Jakob et al, 2011; Tsouroula et al, 2016). HR repair at rDNA loci often results in repeat loss (Warmerdam et al, 2016). In agreement with data derived from cell lines, *in vivo* studies in animal models and meta-analysis of TCGA panel tumors and normal tissue showed that genomic unstable cancers exhibit 45S rDNA repeat loss (Wang & Lemos, 2017; Xu et al, 2017a). The loss of 45S rDNA repeats is considered a consequence of HR break repair of damage that derives from conflicts between transcription and replication in the rapidly dividing cancer cells. RASSF1A is frequently transcriptionally silenced in tumors due to promoter methylation an epigenetic event that associates with early cancer onset in lung cancer and correlates with adverse prognosis in several cancer types (Grawenda & O’Neill, 2015). When we looked for copy number variations of 45S rDNA in a lung adenocarcinoma cohort (LUAD) we found that tumors with high levels of RASSF1A promoter methylation maintain a higher copy number compared to low promoter methylated lung cancers, an observation that suggest compromised HR rDNA repair. We also confirmed maintenance of rDNA repeats in cells in which RASSF1A expression is silenced after exposure to rDNA DSBs. These cells possibly rely on NHEJ for repair that takes place fast after induction of damage in the nucleolar interior (Harding et al, 2015). However, inability to efficiently repair persistent breaks via HR in the absence of the scaffold results in compromised cell viability. Surviving cells potentially harbor extensive genomic instability to which they have probably adapted to, overcoming apoptosis checkpoints. Previous work has shown that RASSF1A can promote p73 mediated apoptosis in response to ATM signaling activation in a Hippo pathway dependent manner, therefore RASSF1A loss could also facilitate escape from apoptosis of genomically unstable cells (Hamilton et al, 2009; Matallanas et al, 2007).

The challenging nature of the nucleoli constitutes a hub of genomic instability. Previous studies have highlighted differences in the regulation of the rDNA damage response compared to other genomic sites including dedicated adaptor proteins (Larsen et al, 2014), differential regulation of ATM/ATR signaling (Korsholm et al, 2019; Mooser et al, 2020), repair of persistent breaks with HR through the cell cycle (van Sluis & McStay, 2015), and specific histone post translational modifications (Pefani et al, 2018). Figuring how the DNA damage response is organized in this area of the genome is important for understanding cancer development and designing novel cancer treatments. Nucleolar transcriptional activity and rDNA copy numbers have been proposed as biomarkers (Warmerdam & Wolthuis, 2019). CX-5461 Pol I inhibitor results in rDNA breaks due to stabilization of R-loops or G-quadruplexes and recombination deficient cancers showed increased sensitivity to the agent as a monotherapy or in combination with PARP inhibitors (Sanij et al, 2020; Xu et al, 2017b). RASSF1A promoter methylation has been proposed as a biomarker in cancer diagnosis (Dubois et al, 2019). In this study, we provide additional mechanistic insight in rDNA DSB break repair and highlight the RASSF1A scaffold as DNA repair factor effectively recruited at rDNA breaks and important for the establishment of the local response. Recruitment of RASSF1A in a small number of sites other than rDNA DSBs upon exposure to γIR or radiomimetic agents, may also serve to amplify local ATM signal where repair is challenging. 53BP1 has been shown to act in establishment of local ATM signals within heterochromatic repeats (Hansen et al, 2016; Noon et al, 2010), therefore future studies would be interesting to address whether the scaffold could facilitate this 53BP1 function in additional chromatin contexts.

## Acknowledgments

We would like to thank Meng Wang and Bernard Lemos (School of Public Health, Harvard) for providing rDNA copy numbers analysis for the LUAD cohort. We would like to thank Brian Mc Stay (NUI Galway) for providing the V5-I-PpoI, V5-I-PpoI H98A plasmids and rDNA probe, Niels Mailand (Novo Nordisk Foundation Research center, Denmark) for providing the 53BP1 constructs and Gaëlle Legube (CBI Toulouse) for providing the AsiSI inducible (DiVa) cell line. We thank the Advanced Light Microscopy Facility at the University of Patras and the Microscopy Scientific Research Facility at the Department of Oncology at the University of Oxford.

## Author contributions

Conceptualization: DEP. Research design: DEP and EON. Experiments and analysis: DEP, ST, GV, FW, MC and AP. Resources: ZL and VG. Funding: DEP and EON. Writing /original draft preparation: ST, DEP and EON. All authors have read the manuscript.

